# Developmental Influences on Symptom Expression in Antipsychotic-Naïve First-Episode Psychosis

**DOI:** 10.1101/2020.06.19.160093

**Authors:** Miranda Bridgwater, Peter Bachman, Brenden Tervo-Clemmens, Gretchen Haas, Rebecca Hayes, Beatriz Luna, Dean F. Salisbury, Maria Jalbrzikowski

## Abstract

**Introduction:** The neurodevelopmental model of psychosis was established over 30 years ago; however, the developmental influence on psychotic symptom expression – how a person’s age affects clinical presentation in first-episode psychosis – has not been thoroughly investigated.

**Method:** Using generalized additive modeling, which allows for linear and non-linear functional forms of age-related change, we leveraged symptom data from a large sample of antipsychotic-naïve individuals experiencing a first episode of psychosis (N=340, 12-40 years, 1-12 visits), collected at the University of Pittsburgh from 1990-2017. We examined relationships between age and severity of perceptual and non-perceptual positive symptoms and negative symptoms. We also tested for age-associated effects on *change* in positive or negative symptom severity following baseline assessment, and explored the time-varying relationship between perceptual and non-perceptual positive symptoms across adolescent development.

**Results:** In the cross-sectional and longitudinal data, perceptual positive symptoms significantly decreased with increasing age (F=7.0, p=0.0007; q=0.003) while non-perceptual positive symptoms increased with increasing age (F=4.1, *p*=0.01, *q*=0.02). These relationships remained significant when SES, IQ, and illness duration were included as covariates. There were no developmental effects on change in positive or negative symptom severity (all *p*>0.25). Finally, an association between severity of non-perceptual and perceptual symptoms developed with increasing age, with a significant association starting at age 18.

**Conclusion:** These findings suggest that as cognitive maturation proceeds, perceptual symptoms attenuate while non-perceptual symptoms are enhanced, reflecting influences of developmental processes on psychosis expression. Findings underscore how pathological brain-behavior relationships vary as a function of development.

## Introduction

Over the past 30 years, the neurodevelopmental model of schizophrenia has become a dominant theoretical framework for organizing findings and generating hypotheses related to psychosis pathogenesis. The premise of the model is that an individual’s sensitivity to certain inputs (e.g., teratogens, perinatal complications, adverse childhood experiences) and likelihood of expressing certain clinically significant outputs (e.g., disorganized behavior, hallucinations) are modulated by the individual’s brain maturation, particularly during adolescence (Weinberger *et al*., 1986; Murray and Lewis, 1987; Insel, 2010; Owen *et al*., 2011; Rapoport *et al*., 2012). Despite the prominence of this model, differences in symptom expression as a result of these maturational changes throughout adolescent and young-adult development have not been examined thoroughly.

In line with the proposed model, late adolescence and early adulthood is a time of increased vulnerability for the emergence of symptoms that meet criteria for schizophrenia-spectrum disorders (Amminger *et al*., 2006; Öngür *et al*., 2009). However, there is not conclusive evidence of symptomatology changing over the course of development in psychosis. If brain maturation modulates the expression of psychosis (both prevalence and severity of symptoms), it is reasonable to expect, for example, that a 12-year old’s symptom expression differs from a 26-year old’s. Symptom expression of other psychiatric disorders, including depression and anxiety, changes across development, particularly during adolescence (DuBois *et al*., 1995; Hankin *et al*., 1998; Garber *et al*., 2002; Van Oort *et al*., 2009; Essau *et al*., 2010); thus, psychotic symptoms could follow a similar pattern. Understanding whether and how age varies with symptom expression could have important implications for creating developmentally-informed assessment and treatment practices, and for our understanding of the mechanisms underlying specific symptoms.

Previous work suggests that age plays an important role in psychosis symptom development. When positive symptoms are divided into specific sub-groups, there is evidence that perceptual positive symptoms (i.e., illusory sensory experiences such as hallucinations) are present to a greater extent in younger individuals (Mueser *et al*., 1990), while non-perceptual positive symptoms (e.g., delusions) have greater prevalence in older individuals with psychosis (Häfner *et al*., 1993). Studies of childhood- or adolescent-onset psychosis find that that these youth endorse higher rates of visual hallucinations than would be expected based on the adult-onset psychosis literature (Green *et al*., 1992; David *et al*., 2011). Furthermore, multiple cross-sectional studies of general population cohorts and individuals at high risk for developing psychosis report that younger individuals are more likely to endorse perceptual psychotic experiences in comparison to older individuals (Kelleher *et al*., 2012b; Brandizzi *et al*., 2014; Schimmelmann *et al*., 2015; Schultze-Lutter *et al*., 2017). However, investigations of age effects on *total* positive symptoms in chronic and first-episode psychosis fail to find differences between age groups or find significant effects of age on symptom presentation (Haas and Sweeney, 1992; Sharma, 1999; Ballageer *et al*., 2005; White *et al*., 2006; Joa *et al*., 2009). Taken together, these results suggest positive symptoms of psychosis display significant age-related variability, but it is critical to examine developmental patterns within relevant sub-groups of positive symptoms. Nonetheless, age effects have not yet been systematically examined in a longitudinal first-episode psychosis sample, which is less likely to be influenced by disease chronicity and medication effects.

Examinations of developmental influence on negative symptom presentation have not had the same level of focus. Some studies have reported that younger people in the early course of a schizophrenia-spectrum disorder showed more prominent negative symptoms (Ballageer *et al*., 2005; Pencer *et al*., 2005), but others have not found this to be the case, particularly for total negative symptoms (Haas and Sweeney, 1992; White *et al*., 2006; Joa *et al*., 2009). Given that late adolescence and early adulthood are periods of dramatic change, both in terms of neurodevelopment and transition to new roles (e.g. starting college, full-time work, etc.), developmentally-focused explorations of negative symptom expression over time may be particularly important as negative symptoms are thought to contribute more to functional impairments than positive symptoms (Ho *et al*., 1998; Milev *et al*., 2005) and cognitive, social, and emotional systems are actively maturing through adolescence (*Pfeifer et al*., 2012; *Luna et al*., 2015).

Table 1 contains a summary of studies investigating age effects in symptom presentation across the psychosis spectrum. While there are some patterns observed in previous work (as described above), there are also inconsistencies. One explanation for these inconsistencies may be that antipsychotic medication exposure obscures age-symptom associations in first-episode samples. All previous studies that report on age effects in those diagnosed with psychotic disorders include individuals who were taking (or had previously taken) antipsychotic medications. Additionally, the majority of the studies are cross-sectional in nature, precluding the ability to assess within-subject change, and the statistical methods used in these studies only assessed linear relationships. Many developmental processes follow a non-linear trajectory and non-linear modeling approaches in developmental neurocognitive science have identified distinct periods of continued refinement of brain structure in typically-developing youth (Simmonds *et al*., 2014; Calabro *et al*., 2020). Use of these approaches with longitudinal symptom data may identify distinct periods of change that are obscured in cross-sectional or linear models. Finally, given evidence that neurobiological factors exert differential influences on symptomatology at distinct points in development (Glaser *et al*., 2011; Jalbrzikowski *et al*., 2017; Ellwood-Lowe *et al*., 2018), use of time-varying approaches may prove to be informative.

**Table 1.**
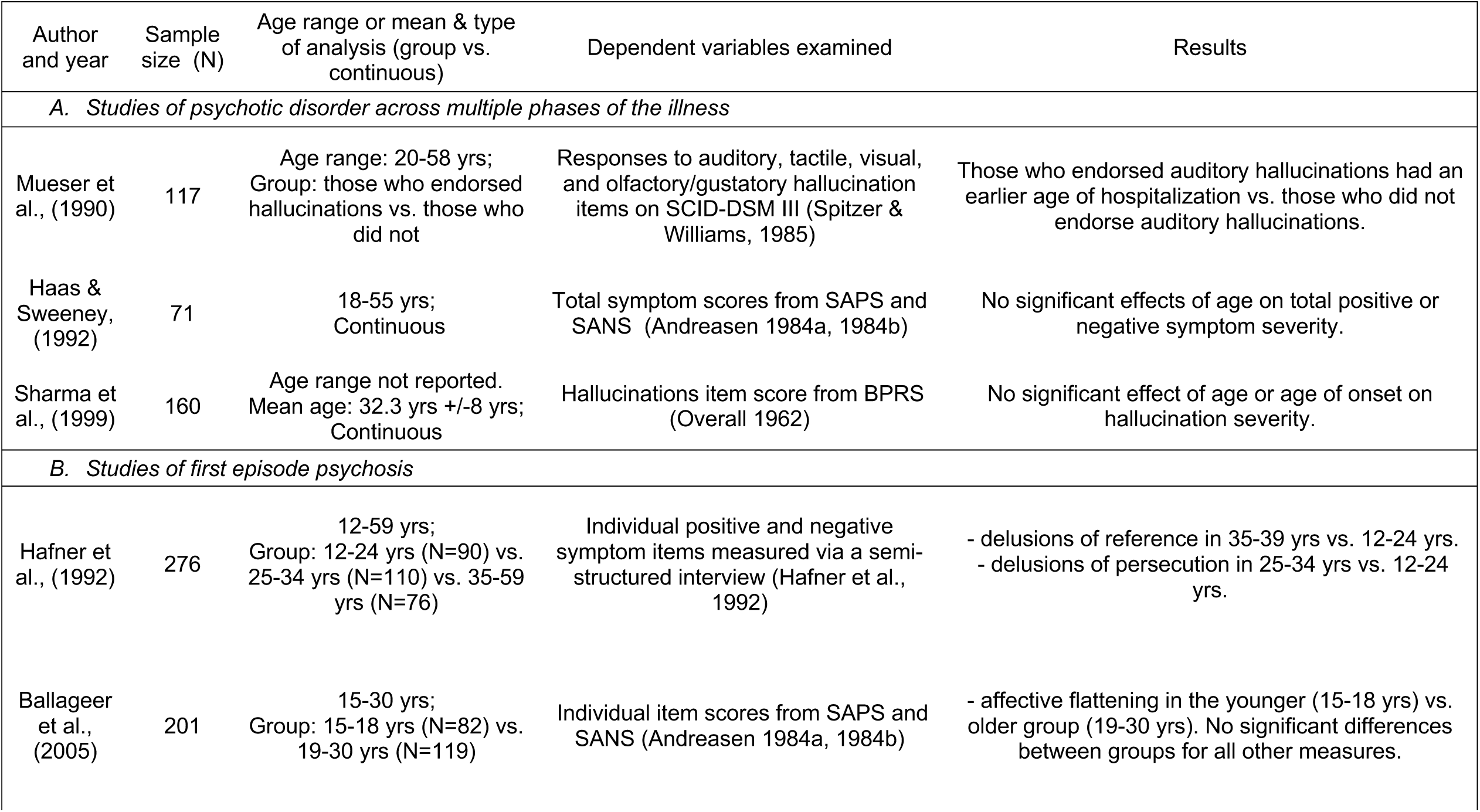
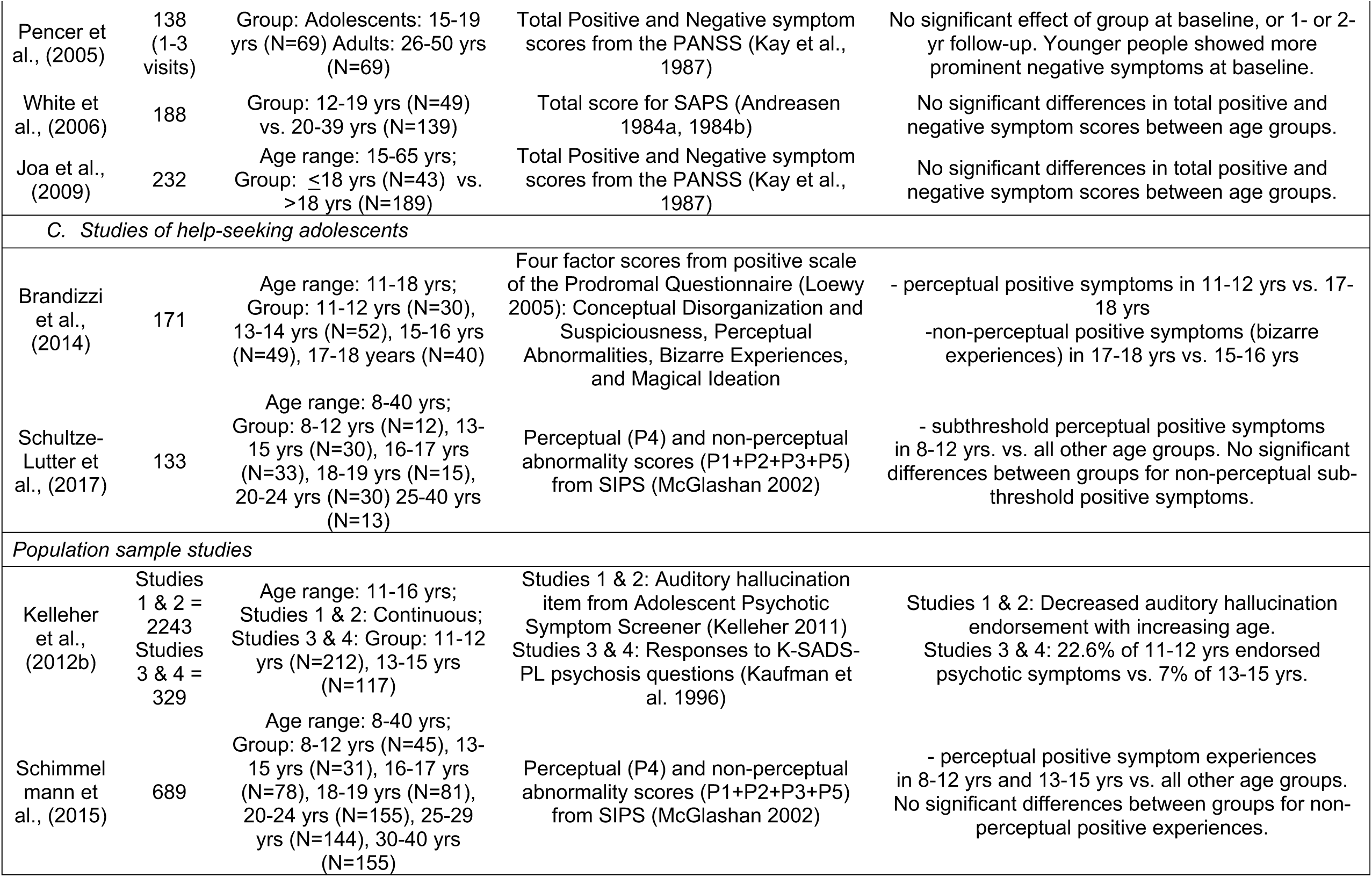
Summary of previous studies that have examined effects of age on psychotic symptom severity. The table is broken down into A) studies that examine participants across the phase of illness, B) studies that focus on first episode psychosis, C) studies of help-seeking adolescents, and D) population sample studies. Other than Pencer et al. (2005), all studies are cross-sectional in nature. All studies of participants with diagnosed with a psychotic disorder include individuals who are currently or previously have been prescribed antipsychotic medication. Abbreviations: yrs.: years; SCID-DSMII= Structured Clinical Interview for the DSM-III; SAPS=Scale for the Assessment of Positive Symptoms; SANS=Scale for the Assessment of Negative Symptoms; BPRS=Brief Psychiatric Rating Scale; PANSS=Positive and Negative Syndrome Scale; SIPS=Structured Interview for Prodromal Syndromes; P1=unusual thoughts rating on SIPS; P2=suspiciousness rating on SIPS; P3=grandiosity rating on SIPS; P4=perceptual abnormality rating on SIPS; P5=disorganized communication on SIPS; KSADS-PL=Kiddie Schedule for Affective Disorders and Schizophrenia, Present and Lifetime version.

In this study, we leveraged a longitudinal sample of antipsychotic-naïve (at baseline) first-episode psychosis participants (FEP, N=340, 1-12 visits, 12-40 years) to 1) examine developmental patterns effects of perceptual and non-perceptual positive symptoms, and negative symptoms, 2) investigate developmental effects on change in psychotic symptom expression following first-episode, and 3) explore age-varying relationships between perceptual and non-perceptual positive symptoms expression. Based on previously reported age-related variance in symptomatology from cross-sectional research, we hypothesized that perceptual positive symptoms would decrease with increasing age, non-perceptual positive symptoms would be stable across adolescent development. All remaining analyses were exploratory.

## Materials and Methods

### Participants

Participant data was taken from archival and ongoing studies at the University of Pittsburgh (1990-2017). The final sample consisted of 340 individuals. See Figure 1 for demographic information and sample characterization. Study procedures were approved by the University of Pittsburgh Institutional Review Board and performed in accordance with the Declaration of Helsinki. All participants or their legal guardians provided written informed consent after study procedures were fully explained.

**Figure 1.**
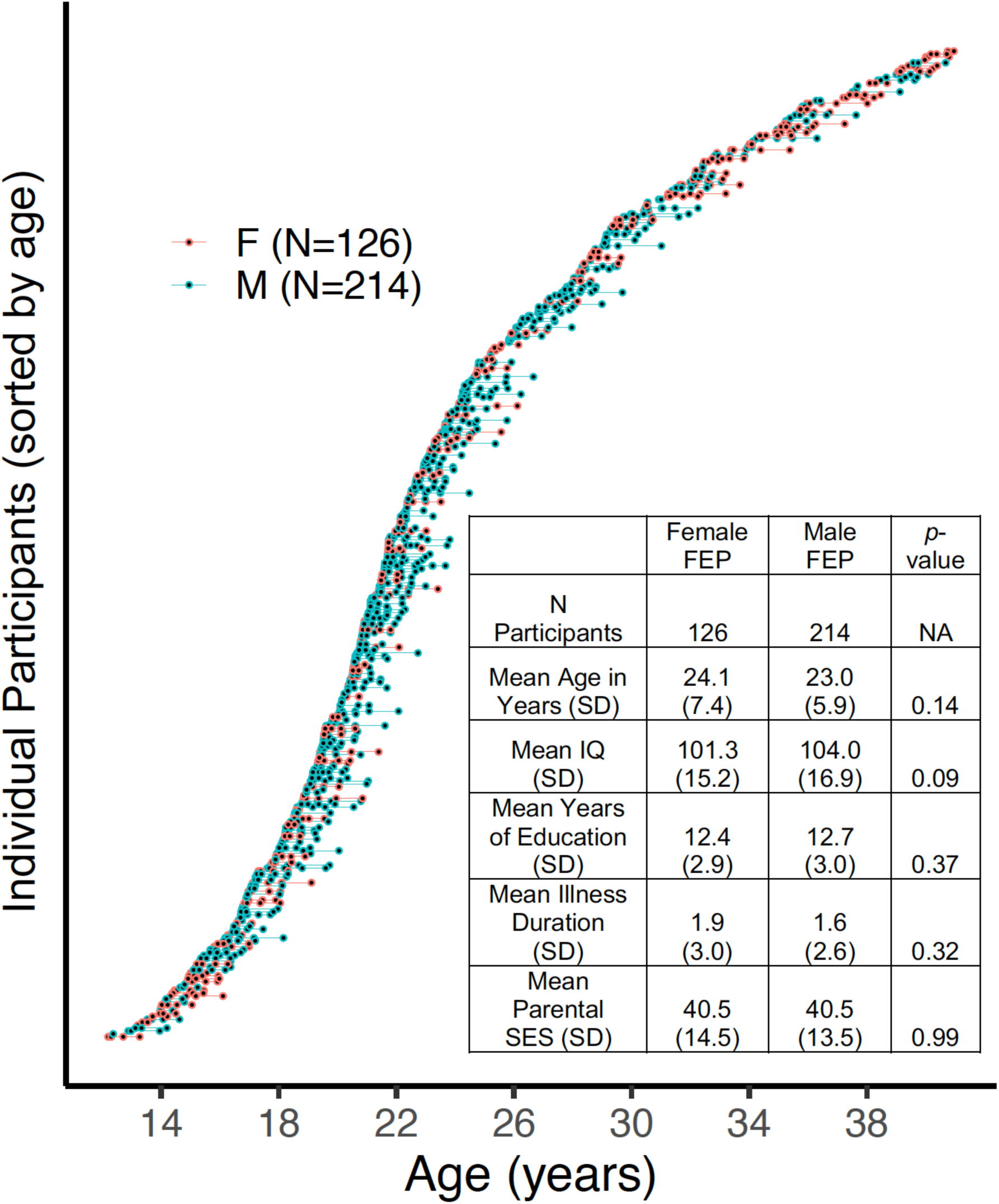
Waterfall plot of all participants and their respective visits (blue circles=male, red circles=female). Each individual circle represents a participant at a particular visit. Lines connecting the circles refer to the time in between visits. A demographic table is in the bottom right of the plot. Two-hundred ninety individuals (85%) had 2 or more visits.

Exclusion criteria for all participants included: medical illness affecting central nervous system function or IQ lower than 75 (as determined using the Wechsler Abbreviated Scale of Intelligence, Wechsler, 1999). Inclusion criteria for FEP were as follows: experiencing one’s first psychotic episode, no specialized prior treatment for his/her psychotic symptoms, and antipsychotic-naive. First-episode psychosis diagnoses were determined using all available clinical information and data gathered from a Structured Clinical Interview for DSM-IV (SCID, First *et al*., 2002) conducted by a trained clinician. Senior diagnostician/clinical researchers confirmed diagnoses at consensus meetings. Illness duration for each client was also determined in the consensus conference after a review of historical information about psychosis onset. See Supplementary Figure S1 for a detailed description of participants removed from final analyses.

### Clinical Measures

We assessed positive symptom severity with the Scale for the Assessment of Positive Symptoms (SAPS; Andreasen, 1984b). The SAPS includes 34 items addressing hallucinations, delusions, bizarre behavior and formal thought disorder on a 0 (absent) to 5 (severe) scale. Consistent with Schimmelmann et al.(2015), we summed individual items from the SAPS (not including the SAPS global rating items) to calculate perceptual (SAPS items 1-6, range: 0-30) and non-perceptual (SAPS items 8-33, range: 0-120) positive symptom scores.

We assessed negative symptom severity with the Scale for the Assessment of Negative Symptoms (SANS; Andreasen, 1984a). The SANS includes 25 items addressing affective flattening, alogia, avolition, anhedonia and attention on a 1 (absent/mild) to 5 (severe) scale (range: 25-125). All 25 items were initially scored for a total negative symptom score (not including global rating items) and were then broken down into respective subgroups.

### Statistical Analyses

#### Aim 1: Developmental effects of symptomatology in FEP

To assess developmental effects of symptomatology in FEP, data were modeled using penalized splines within a general additive model (GAM; Hastie and Tibshirani, 1986, 1990; Wood, 2017). A GAM is an extension of the general linear model that does not assume that the relationship between independent and dependent variables is linear, allowing for a more flexible predictor. Smoothed predictor function(s) are automatically derived *during* model estimation through the use of basis functions (here, thin plate splines: MCGV default). Because incorporation of more basis functions incurs greater penalties (using restricted maximum likelihood), GAM is able to address many limitations of other non-linear models (e.g., over-fitting, variance/bias trade-offs). The dependent variable was the respective clinical measure being assessed. Fixed effects entered into the model were baseline age, visit, and sex. Subject was included as a random effect (*r*), allowing us to model and account for the non-independence of longitudinal data (multiple visits). Because all clinical symptom data was skewed to the left, we performed a log transformation to normalize symptom distributions.

Because there are known sex differences in psychosis age of onset, we first explored smoothed effects for age in males and females separately by including a moderating effect of sex on age:

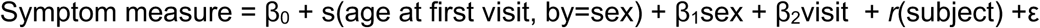

We also tested smoothed age effects of the model for both sexes aggregated together:

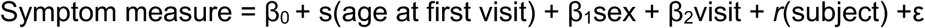

To determine the best model fit from the two above models, we used Bayesian information criterion (BIC), a commonly used measure for model selection (Vrieze, 2012).

The broad age range and longitudinal data structure of the study (see Figure 1) allowed us to explore both a) developmental effects of symptom expression at first episode and effects of b) illness chronicity. To explore these potentially diverging developmental effects, we first included baseline age and visit as separate predictors in the GAM. However, despite having these entered as separate regressors, by including longitudinal data, we could hypothetically fail to truly measure symptoms at first expression. Thus, we re-ran all analyses on the cross-sectional data only. We also tested the effects of socioeconomic status, cognition, and illness duration on the above models.

In order to identify specific developmental periods that may have significant age-related change in symptom expression, we performed a posterior simulation on the first derivative of GAM fits. As in recent work from our group (Calabro *et al*., 2020) and following established guidelines (Wood, 2017), we used a multivariate normal distribution whose vector of means and covariance were defined by the fitted GAM parameters to simulate 10,000 GAM fits and their first derivatives (generated at 0.1 year age intervals). Significant intervals of age-related change in symptom expression were defined as those ages when the confidence intervals (95%) of these simulated GAM fits did not include zero (p < .05).

#### Aim 2: Developmental effects of change in positive and negative symptom expression in FEP

To assess developmental effects of change in symptom measures, we selected the first two visits from participants with multiple time points (N= 290), which ranged between 0-24 months (Supplementary Figure S2). This maximized sample size, as two visits was the most common type of longitudinal data. We then created change scores for each symptom measure (symptom measure at visit 2-symptom measure at visit 1). We again used GAM and modeled the change score as the dependent variable and the smoothed effect of age as the predictor. We controlled for symptom expression at the first visit in the model, to account for regression to the mean and initial level of symptom severity. Similar to the analyses above, we used BIC to determine the best model fit with respect to sex and then assessed the smoothed effects of age:

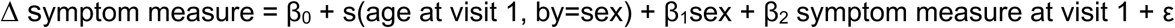

vs.

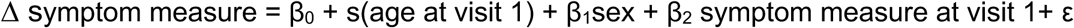

We also re-ran these analyses and included the number of days between visit 1 and visit 2 as a covariate.

Aim 3: Interaction between non-perceptual positive symptoms and age on perceptual positive symptoms

We tested how the smoothed effect of age on perceptual positive symptoms varies according the degree of non-perceptual positive symptoms; i.e., the effect of a smoothed interaction between age and non-perceptual positive symptoms:

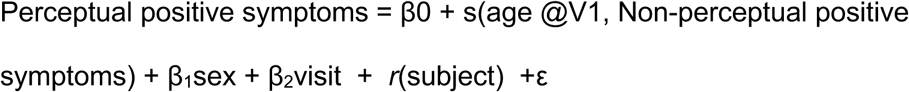

We used contour plots (using mcgv package in R; Wood, 2011) to visualize the result. We also explored the interactions between negative symptoms and age on total positive symptoms, as well as perceptual and non-perceptual positive symptoms.

Within each set of the above analyses (Aims 1-3), false discovery rate (FDR) was used to correct for multiple comparisons (*q*<.05)

## Results

### The presentation of particular positive symptoms changes across adolescent development

Table 2 reports the results of the smoothed effect of age on psychotic symptom expression. Perceptual positive symptoms expressed at first episode declined with increasing age, longitudinally (*F*=7.0, *p*= 7.0e-04; *q*=0.003, Figure 2A). Significant periods of age-related changes occurred between 14.3 and 26.8 years of age. These age effects were driven by auditory and visual hallucinations, while developmental trajectories for somatic and olfactory hallucinations remained stable from 12-40 years (Supplementary Figure S3). Non-perceptual positive symptoms at first episode significantly increased with increasing age (*F*=4.1, *p*=0.01, *q*=0.02, Figure 2B). Significant periods of age-related changes occurred between 16.3-22.4 years old. These age-associated increases were driven by delusions and thought disorder (Supplementary Figure S4).

**Table 2.**
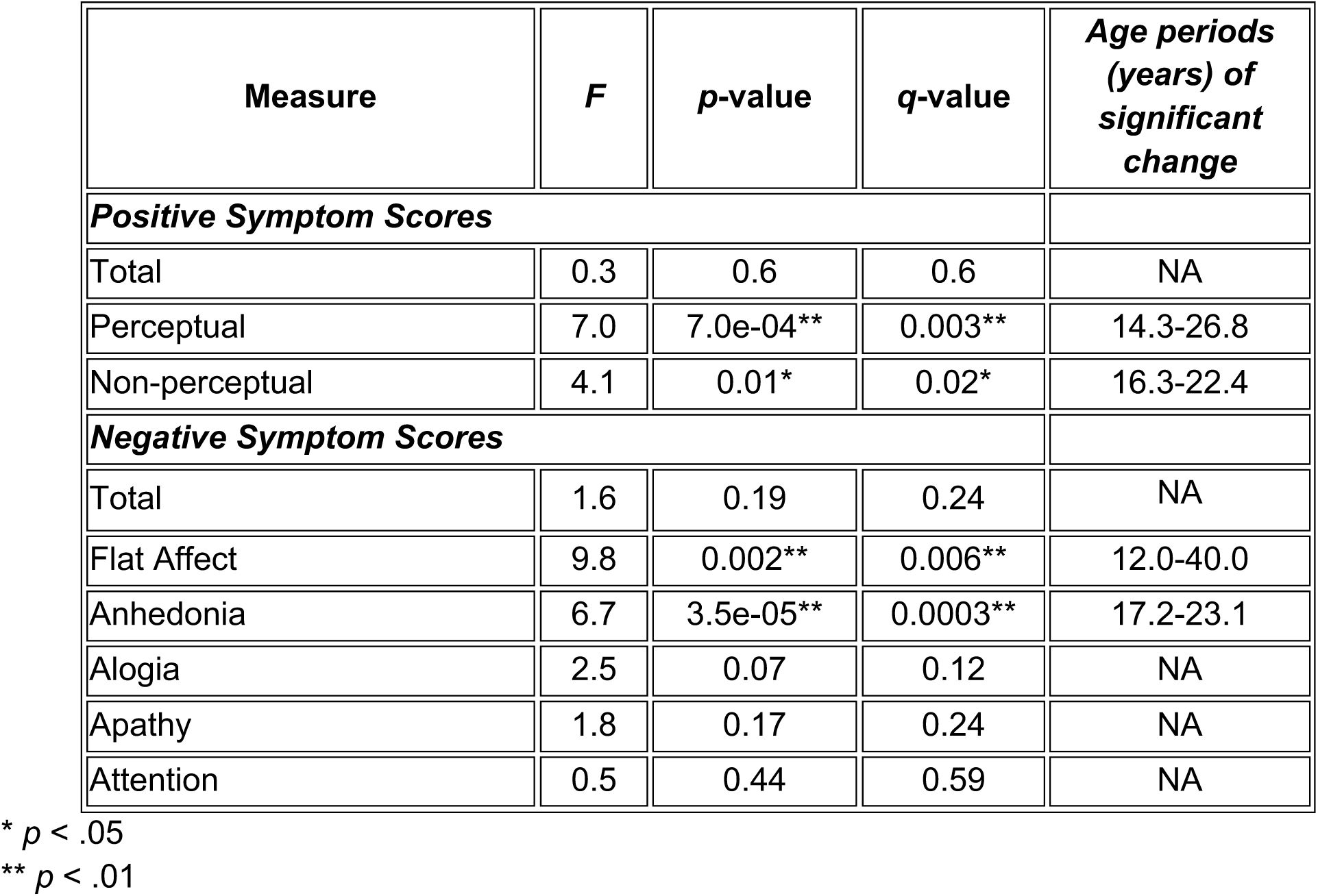
Developmental effects on positive and negative symptoms in first episode psychosis.

**Figure 2.**
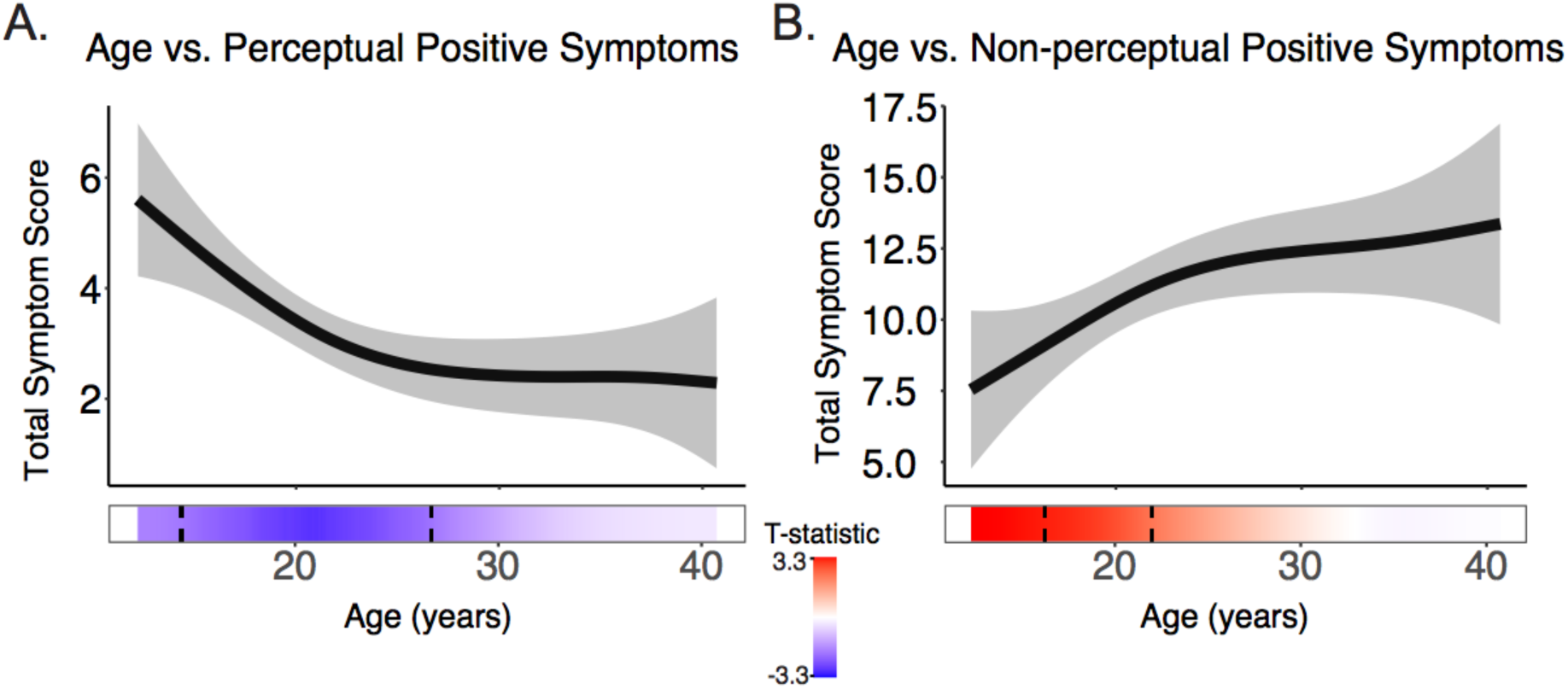
A) Perceptual positive symptoms significantly decreased with increasing age longitudinally, while B) non-perceptual positive symptoms increased with increasing age. The bar underneath the age plot reflects the derivative of the slope, i.e., the rates of change taking place at a particular age, scaled as a pseudo t-statistic, based on the posterior simulation. The dotted lines indicate when significant age associated change is taking place. Brighter red indicates greater age-related increases, while bright blue indicated greater age-related decreases.

There was not a significant effect of age on total positive symptoms longitudinally (*F*=0.3, *p*=0.6, *q*=0.6). For all models tested, there were no significant main effects of sex (all *p*>0.5). Furthermore, for all models, BIC estimates showed that including the effect of sex on smoothed age (i.e., sex by smoothed age interaction) did not significantly improve model fit (Supplementary Table S1).

All age-related changes remained statistically significant (*p*<.05) when intelligence quotient, average parental socioeconomic status, and illness duration were included in the model (Supplementary Tables S2-S4). Despite changes in how age was modeled, results remained consistent when only cross-sectional data (baseline only) was included in the analyses and when age at each visit (instead of age at visit 1) was used in the analyses (Supplementary Tables S5-S6).

### Exploratory developmental effects of negative symptoms

The level of overall negative symptoms did not change across adolescent development (*F*=1.6, *p*=0.19, *q*=0.24). However, when individual symptoms were examined, presentation of anhedonia increased with increasing age (*F*=6.7, p=3.5e-05, q=0.0003), while flat affect decreased with increasing age (*F*=9.8, *p*=0.002, *q*=0.006). Symptom expression of alogia, attention and apathy remained stable from ages 12-40 years old. Results are presented in Supplementary Table S7 and Figure S5.

### No significant effects of development on symptom change following baseline assessment

There were no significant developmental effects on change in post-baseline symptom expression for any symptom measure (symptom measure at visit 2-symptom measure at visit 1; Figure 3; Supplementary Table S8). The mean length of time between visits was 73.3 days, though there was wide variability in amount of time between assessments (+/-73.5 days, range: 6-700 days). On average, symptom severity was lower in the second visit compared to the first visit, regardless of age (all *p*-values > 0.20). When the number of days between the two visits was included as a covariate, the results remained consistent.

**Figure 3.**
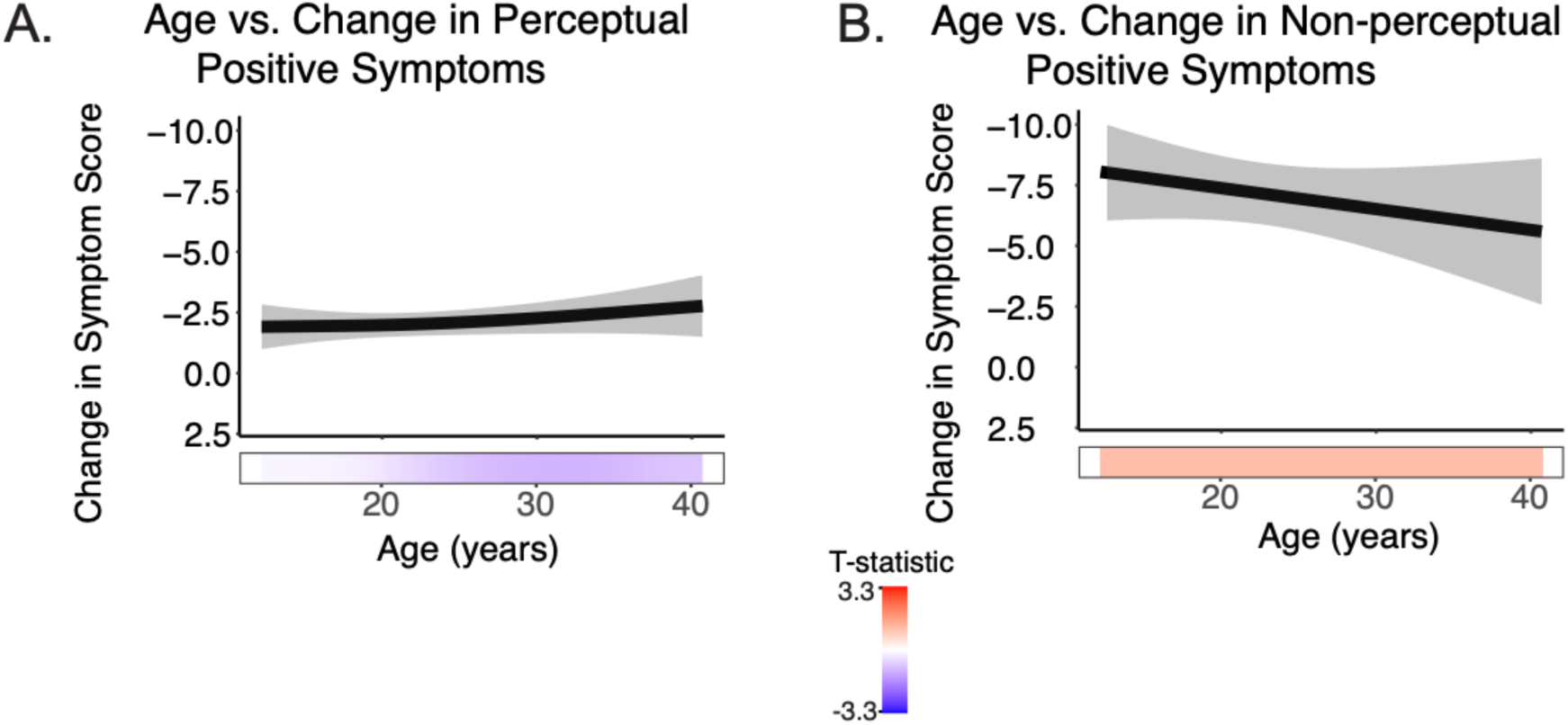
Change in symptom expression remained stable across age for A) perceptual positive symptoms and B) non-perceptual positive symptoms.

### Smoothed Interaction Between Age and Non-Perceptual Positive Symptoms on Perceptual Positive Symptoms

There was a statistically significant interaction between the smoothed effect of age and perceptual positive symptoms on non-perceptual positive symptoms across development (*F*=13.1, *p*=2e-16, Figure 4A). Specifically, in youth (< 18 years), there was ***not*** a statistically significant relationship between perceptual symptoms and non-perceptual symptoms (<18 years, *b*=0.18, *p*=0.11, Figure 4B). However, in adults (≥ 18 years), there was a statistically significant relationship, as higher levels of perceptual positive symptoms were associated with greater levels of non-perceptual symptoms(18-29 years: b=0.38, p=3.2e-11, Figure 4C; 30-40 years: b=0.60, p=1.2e-8, Figure 4D).

**Figure 4.**
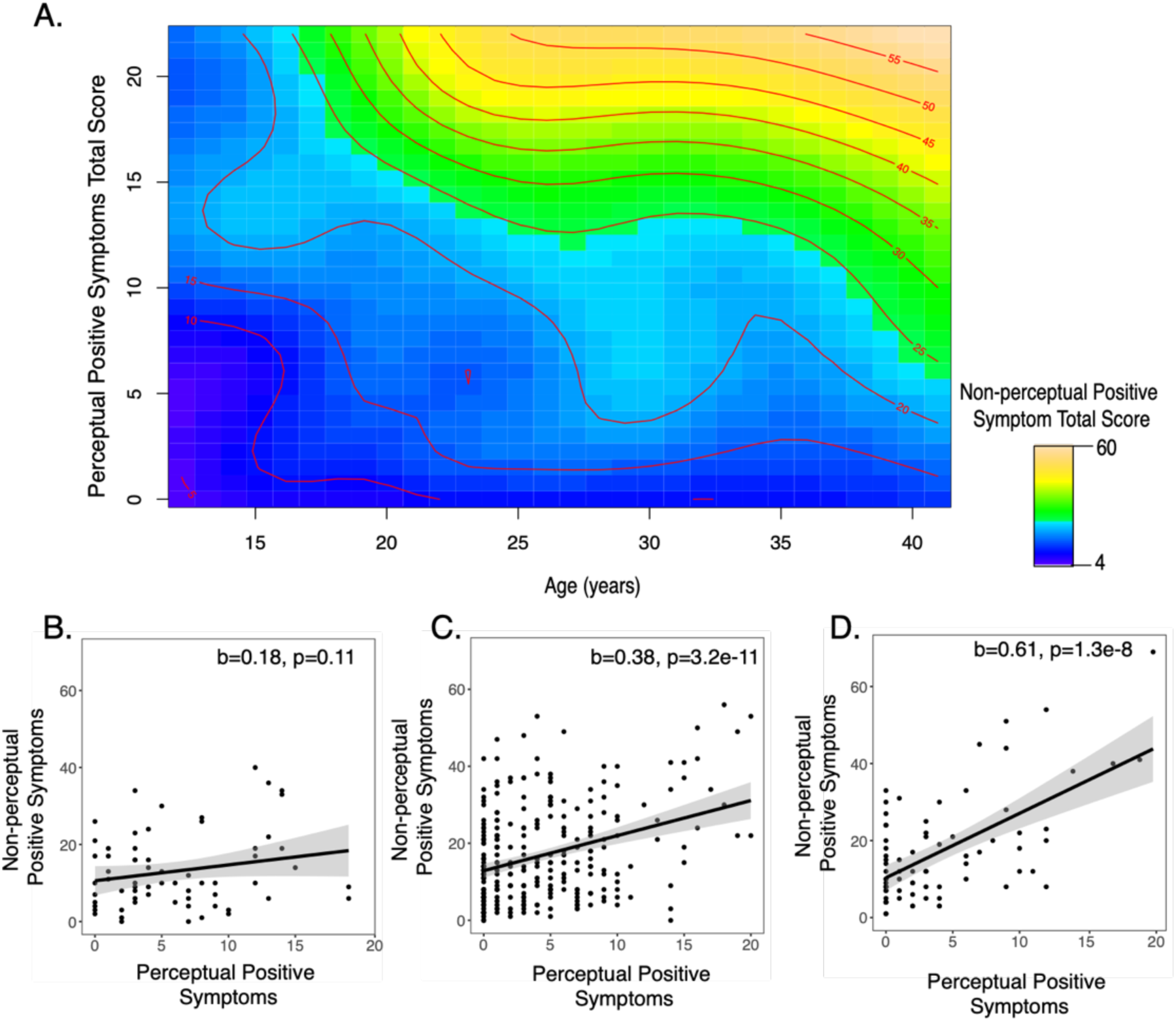
A) A contour plot illustrating how the relationship between perceptual and non-perceptual positive symptoms changes across adolescent development. The color reflects the strength of the severity of non-perceptual positive symptoms, with yellows indicating a higher level of non-perceptual positive symptoms. The severity level of non-perceptual symptoms across different ages is also indicated with red lines and text. To understand this figure, it is helpful to pick a particular age and traverse the height of the graph. At age 15, individuals with greater levels of perceptual positive symptoms (e.g., a score > 10) may have a limited range in the severity of non-perceptual positive symptoms (15-20) and the variables are not strongly associated with one another. At age 35, as individuals’ levels of perceptual positive symptoms increase, their non-perceptual positive symptoms also increase (the change from blue to yellow, and the successive increase in non-perceptual positive symptom severity, observed by the multiple red lines on the right-hand side of the graph). For visualization purposes, we also plot the linear fit between perceptual and non-perceptual positive symptoms in three separate age ranges: B) 12-17.9 years old, C) 18-29.9 years old, and D) 30-40 years old.

## Discussion

By examining developmental changes in symptom expression in a large, antipsychotic-naïve sample of individuals experiencing their first episode of psychosis (12-40 years old), we found distinct patterns of association between development and particular psychotic symptoms. Neither medication exposure at follow-up assessments, chronicity, intelligence, nor socioeconomic status accounted for these associations. We therefore consider these results evidence of selective age-related developmental influences on emerging psychosis – a key tenet of the neurodevelopmental model. Additionally, the nature of the age-symptom associations may inform our understanding of the pathophysiological processes underlying first-episode psychosis, highlighting the importance of developmentally informed approaches for both research and treatment in this population.

### Distinct developmental trajectories of specific positive symptom expression

We used nonlinear modeling strategies to determine distinct periods of change in positive symptom expression. Specifically, between 14-26 years old, perceptual symptom severity decreased significantly, particularly for auditory and visual hallucinations. These findings are consistent with reports that hallucinations occur in higher percentages of cases of childhood-and adolescent-onset psychosis, as compared to adult-onset psychosis (Green *et al*., 1992; David *et al*., 2011) and dovetail nicely with clinically-ascertained high-risk and population sample findings that younger adolescents are more likely to report perceptual abnormalities than older adolescents and young adults (Kelleher *et al*., 2012b, 2012a; Brandizzi *et al*., 2014; Schimmelmann *et al*., 2015; Schultze-Lutter *et al*., 2017). Taken together, these findings suggest that across the continuum of psychosis-spectrum severity (i.e. ranging in severity from psychotic-like experiences to diagnosable psychotic disorders), a decrease in perceptual positive experiences occurs with increasing age and may reflect the period of specialization that is indicative of adolescent development.

In contrast, non-perceptual positive symptom expression significantly increased with increasing age from 16-22 years old, an effect driven by delusions and thought disorder. These findings closely align with those of Hafner et al., (1992), who showed that, in a chronic schizophrenia-spectrum sample, older participants (>25 years) were more likely to endorse delusions in comparison to younger participants (ages 12-24 years). Together, these findings suggest that delusions may be less severe or likely to form in early adolescence, or that they are less impairing or distressing (therefore less likely to be reported to clinicians). Longitudinal investigations of the development of non-perceptual positive symptoms (specifically, delusions and formal thought disorder) and functional impairment are needed to clarify how young adults experiencing psychosis are affected functionally.

These developmental differences in perceptual and non-perceptual symptom expression point to potentially distinct treatment needs for individuals diagnosed with psychosis-spectrum disorders in childhood or adolescence rather than adulthood. For example, clients in early adolescence may benefit from learning strategies that target effective ways to respond to hallucinations, whereas, in older clients, it may be more effective to focus on cognitive reappraisal to cope with delusional thoughts. Alternatively, the developmental variations we observed could reflect the fact that symptom expression has different clinical implications at different ages. For example, it is well-documented that types of stressors change across adolescent development (Compas, 1987; Simmons *et al*., 1987; Eccles *et al*., 1993; Stroud *et al*., 2009); perhaps perceptual symptoms are likely to present themselves with stressors that are typical of late childhood/early adolescence, while non-perceptual symptoms are a response to adult stressors. Another possibility is that the developmental timing of a particular risk factor (e.g., trauma, substance use, social adversity) may bring about different types of symptom responses, a phenomenon observed in other psychiatric disorders (see Thapar and Riglin, 2020 for a more thorough discussion).

### Distinct developmental trajectories of specific negative symptom expression

Among negative symptoms, affective flattening expression exhibited consistent linear decreases with increasing age, while anhedonia severity increased with increasing age between 17 and 23 years. There were no significant age-related changes found for alogia, attention or apathy. Our findings of decreased affective flattening with increasing age are consistent with previous work (Ballageer et al 2005, Hafner et al. 1992). However, the majority of the previous literature has examined *overall* negative symptom severity rather than changes in individual negative symptoms; and these studies have reported no significant differences in overall negative symptom severity by age of onset (Haas and Sweeney, 1992; White *et al*., 2006; Joa *et al*., 2009) -findings which are replicated in this study. Further work examining age effects on individual negative symptoms, as opposed to overall negative symptom severity, should be done. For example, while increased severity of negative symptoms has been repeatedly linked to greater functional impairment and lower quality of life (Ho *et al*., 1998; Herbener and Harrow, 2004; Mäkinen *et al*., 2008; Ventura *et al*., 2009; Fulford *et al*., 2013; Santesteban-Echarri *et al*., 2017), it is unknown if this relationship is stable across adolescent development, and to what extent specific negative symptoms contribute to this association. For instance, in adolescence, affective blunting may contribute to functional impairment by making peer social interactions more difficult, while worsening social anhedonia/avolition may contribute to (or represent) functional impairment later in life.

### No evidence of developmental effects of change in positive and negative symptom expression

There were no significant developmental effects on *change* in symptom expression across study visits. Across our age range, symptom severity was significantly lower at the second visit. An earlier onset of psychosis (before age 18) is generally considered to be worse for long-term outcome (Clemmensen *et al*., 2012; Immonen *et al*., 2017). However, within a shorter time frame (mean length of time between assessments: ∼3 months), our results suggest that change in symptom severity is similar across development. Others have found that earlier age of onset is associated with increased time to symptom remission in first-episode samples (Malla *et al*., 2006; Veru *et al*., 2016). Unlike our study, these studies did not quantitatively assess change in symptom presentation (regardless of direction). Furthermore, these studies did not focus specifically on medication-naïve cases; thus, symptom severity may have been associated, in part, with duration of exposure to medication prior to baseline assessment.

### Significant interaction between effect of age and perceptual positive symptoms on non-perceptual positive symptoms

We found that, with increasing age, the relationship between severity of perceptual positive symptoms and non-perceptual positive symptoms grows significantly stronger. Different types of positive symptoms may exhibit distinct functional influence on other types of positive symptoms across development. For example, worsening delusions may help to crystallize or solidify the content, severity, or attribution of perceptual abnormalities – but only in more mature individuals. Abstract and relational reasoning continues to mature during adolescence (Marini and Case, 1994; Crone *et al*., 2009), and older individuals experiencing their first episode of psychosis may use these increased capabilities to help them make sense of perceptual positive symptom experiences they are having. Alternatively, the underlying factor structure of the set of positive symptoms differs between relatively younger adolescents, older adolescents and adults. For example, among younger individuals, perceptual abnormalities load more strongly on a general psychopathology factor (Kelleher *et al*., 2012b; Lancefield *et al*., 2016), while among relatively older individuals, the emergence of perceptual abnormalities may reflect a more specific pathology (i.e. psychosis-spectrum disorders).

### Possible mechanisms underlying developmental changes in symptom expression

The physiological underpinnings of any developmental influences likely vary between the different symptoms or types of symptoms in question. Current hypotheses regarding the nature of developmental changes associated with psychosis onset include how changes in the balance of excitatory and inhibitory influences reflect functionally distinct brain circuits (e.g., Mikanmaa *et al*., 2019). Dysregulated development of dopamine may contribute to this excitatory-inhibitory imbalance, potentially leading to the development of psychotic symptoms (Kapur, 2003; van Nimwegen *et al*., 2005; Larsen and Luna, 2018). Given that perceptual and non-perceptual symptoms are associated with alterations in particular brain networks (e.g., abnormal perceptual experiences may reflect abnormalities in sensory and temporal regions, while delusions may be due to disruptions connections between frontolimbic areas, Corlett *et al*., 2010, 2019; Jardri *et al*., 2013), it is possible that age-associated differences in symptom expression may reflect changes in the excitatory-inhibitory balance of distinct areas of the brain. In future work, particular circuits associated with expression of these symptoms should be studied within a developmental framework.

### Limitations

This study is not without limitations. As data was retrospectively collected from multiple studies, there was variation in the amount of time between visits (range 6-700 days). Replication in a longitudinal study that utilizes uniform follow-up visits is necessary. Furthermore, though all participants were antipsychotic-naïve at baseline, the type of treatment participants engaged in post-baseline (i.e. pharmacological, psychosocial, inpatient/outpatient, and nature and dosages of antipsychotic medications at follow-up assessments) was mixed and not controlled for in this study. Thus, treatment exposure may have partially confounded estimation of age effects on initial (visit 1 to visit 2) symptom change, where all participants showed clinical improvements (decreased symptom presentation). Nonetheless, estimation of age-related developmental effects on the nature and severity of initial presenting symptoms of young people in their first episode of psychosis is an important step in investigating developmental underpinnings of early symptom presentation. Finally, pubertal development and hormonal changes have been implicated as factors that impact the risk for and sex-related variation in age at onset of psychosis (e.g., Corcoran *et al*., 2003; Markham *et al*., 2012; Walker and Bollini, 2002; Walker *et al*., 2008); thus, future work should assess how measures of pubertal development relate to positive and negative symptomatology.

### Conclusion and Future Directions

In addition to finding distinct age-related developmental effects on psychotic symptoms in a sample of clients experiencing their first episode of psychosis, these findings point to the importance of age as an index of developmental effects on specific symptom domains rather than overall symptom severity. Future investigation of specific age-related symptom trajectories may be informative for improving identification of risk factors for psychosis. Presently, only ∼20-30% of the individuals identified as being at high risk for psychosis develop a full psychotic disorder (Fusar-Poli *et al*., 2012); thus, approaching risk characterization and prediction from a developmental perspective may improve identification efforts. Studies of clinical-high risk cohorts report that higher levels of non-perceptual positive symptoms (e.g., unusual thought content and suspiciousness (Cannon *et al*., 2008; Cannon *et al*., 2016)) significantly predict conversion to psychosis, not perceptual positive symptoms, highlighting the important potential for development in future studies of psychosis-risk. Finally, to better understand brain mechanisms underlying these developmental effects on symptom expression, it would be useful to conduct a longitudinal neuroimaging study that examines the relationship of these developmentally-divergent symptoms and distinct neural regions involved in perceptual and non-perceptual positive symptoms.

## Acknowledgements

The project described was supported by the National Institutes of Health through grants K01 MH112774 (Maria Jalbrzikowski, PI); R01 MH094328, R01 MH108568P50 (Dean F. Salisbury, PI); MH103204, P50 MH084053, P50 MH045146 (David A. Lewis, MD, Director); UL1 TR001857, UL1 RR024153 (Steven E. Reis, MD, PI); and M01 RR00056 (Arthur Levine, MD, PI).

We thank the faculty and staff of the WPH Psychosis Recruitment and Assessment Core for their assistance in diagnostic and psychopathological assessments. We also thank Leah Vines and Sabrina Catalano for providing feedback to drafts of the manuscript.

---

**STable 1:**
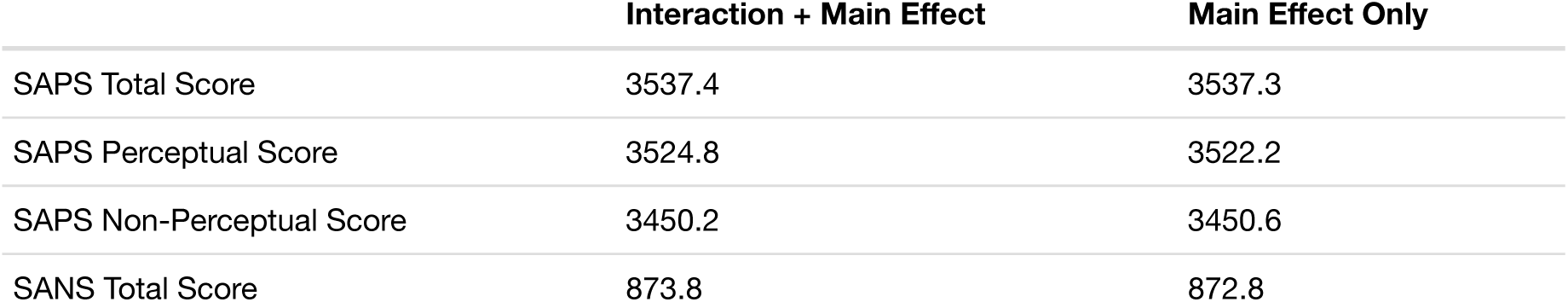
Bayesian Information Criterion estimates of model fit with and without interaction of sex with smoothed age effect (main effect). SAPS=Scale for the Assessment of Positive Symptoms; SANS=Scale for the Assessment of Negative Symptoms

|  | **Interaction + Main Effect** | **Main Effect Only** |
| --- | --- | --- |
| SAPS Total Score | 3537.4 | 3537.3 |
| SAPS Perceptual Score | 3524.8 | 3522.2 |
| SAPS Non-Perceptual Score | 3450.2 | 3450.6 |
| SANS Total Score | 873.8 | 872.8 |

**STable 2:**
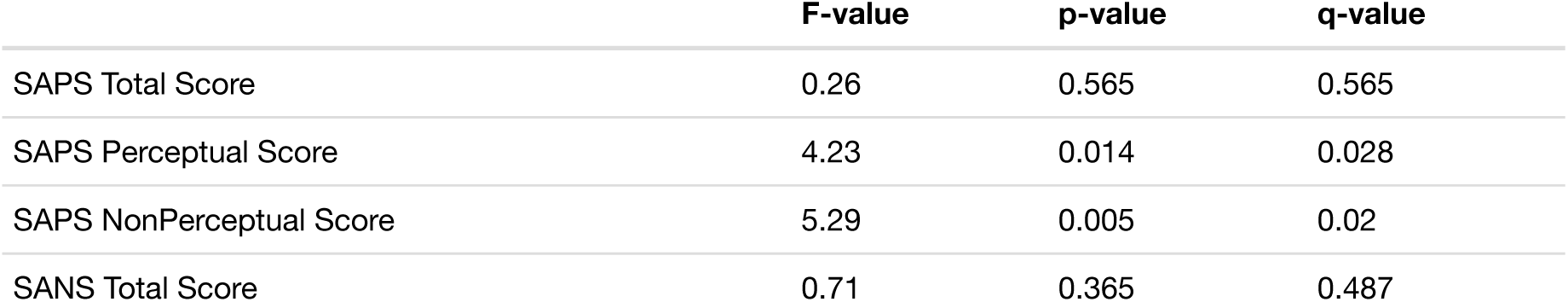
Developmental effects on psychotic symptoms in first episode psychosis when Intelligence Quotient (IQ) is included as covariate.SAPS=Scale for the Assessment of Positive Symptoms; SANS=Scale for the Assessment of Negative Symptoms

|  | **F-value** | **p-value** | **q-value** |
| --- | --- | --- | --- |
| SAPS Total Score | 0.26 | 0.565 | 0.565 |
| SAPS Perceptual Score | 4.23 | 0.014 | 0.028 |
| SAPS NonPerceptual Score | 5.29 | 0.005 | 0.02 |
| SANS Total Score | 0.71 | 0.365 | 0.487 |

**STable 3:** Developmental effects on psychotic symptoms in first episode psychosis when parental socioeconomic status is included as covariate. SAPS=Scale for the Assessment of Positive Symptoms; SANS=Scale for the Assessment of Negative Symptoms

|  | **F-value** | **p-value** | **q-value** |
| --- | --- | --- | --- |
| SAPS Total Score | 0.17 | 0.666 | 0.725 |
| SAPS Perceptual Score | 7.40 | 0.002 | 0.008 |
| SAPS NonPerceptual Score | 2.73 | 0.044 | 0.088 |
| SANS Total Score | 0.12 | 0.725 | 0.725 |

**STable 4:** Developmental effects on psychotic symptoms in first episode psychosis when illness duration is included as covariate in the model. SAPS=Scale for the Assessment of Positive Symptoms; SANS=Scale for the Assessment of Negative Symptoms

|  | **F-value** | **p-value** | **q-value** |
| --- | --- | --- | --- |
| SAPS Total Score | 0.04 | 0.846 | 0.954 |
| SAPS Perceptual Score | 8.32 | 1.93e-04 | 7.72e-04 |
| SAPS NonPerceptual Score | 2.32 | 0.065 | 0.13 |
| SANS Total Score | 0.00 | 0.954 | 0.954 |

**STable 5:** Age-associated effects of psychotic symptoms in first episode psychosis when only cross-sectional data (visit 1) is included in the model.SAPS=Scale for the Assessment of Positive Symptoms; SANS=Scale for the Assessment of Negative Symptoms

|  | **F-value** | **p-value** | **q-value** |
| --- | --- | --- | --- |
| SAPS Total Score | 1.64 | 0.201 | 0.268 |
| SAPS Perceptual Score | 4.37 | 0.004 | 0.016 |
| SAPS NonPerceptual Score | 4.62 | 0.009 | 0.018 |
| SANS Total Score | 0.39 | 0.534 | 0.534 |

**STable 6:** Developmental effects on psychotic symptoms in first episode psychosis when age at first visit is included as a main effect for all visits. SAPS=Scale for the Assessment of Positive Symptoms; SANS=Scale for the Assessment of Negative Symptoms

|  | **F-value** | **p-value** | **q-value** |
| --- | --- | --- | --- |
| SAPS Total Score | 0.24 | 0.604 | 0.805 |
| SAPS Perceptual Score | 7.43 | 5.99e-04 | 0.002 |
| SAPS NonPerceptual Score | 4.04 | 0.016 | 0.032 |
| SANS Total Score | 0.00 | 0.952 | 0.952 |

**STable 7:** Developmental effects on negative symptoms in first episode psychosis.SANS=Scale for the Assessment of Negative Symptoms

|  | **F-value** | **p-value** | **q-value** |
| --- | --- | --- | --- |
| SANS Flat Affect Score | 6.39 | 0.002 | 0.005 |
| SANS Anhedonia Score | 7.88 | 3.29e-04 | 0.002 |
| SANS Alogia Score | 2.89 | 0.043 | 0.072 |
| SANS Apathy Score | 0.60 | 0.477 | 0.477 |
| SANS Attention Score | 1.52 | 0.221 | 0.276 |

**STable 8:**
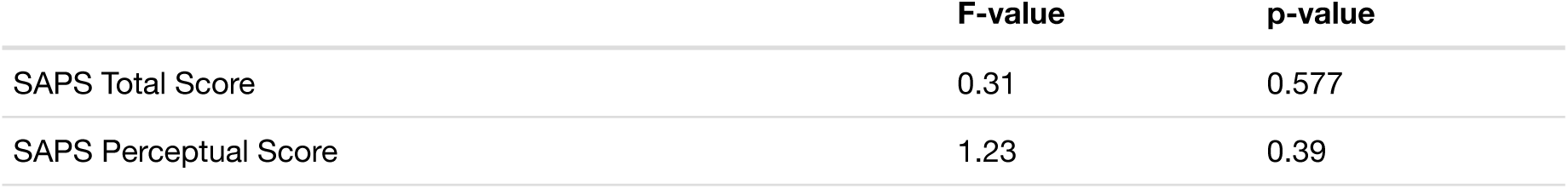
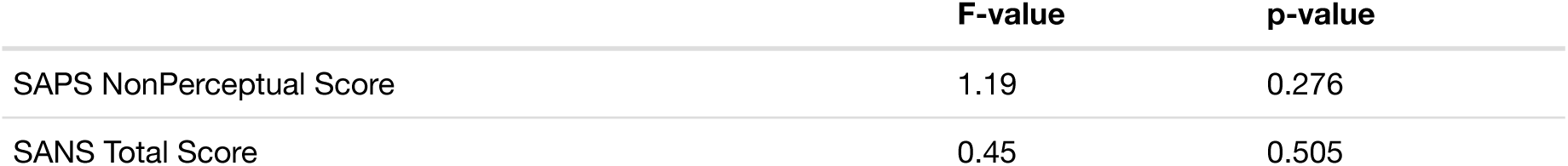
Change in symptom presentation in first episode psychosis (visit 1 - visit 2).

**SFigure 1:**
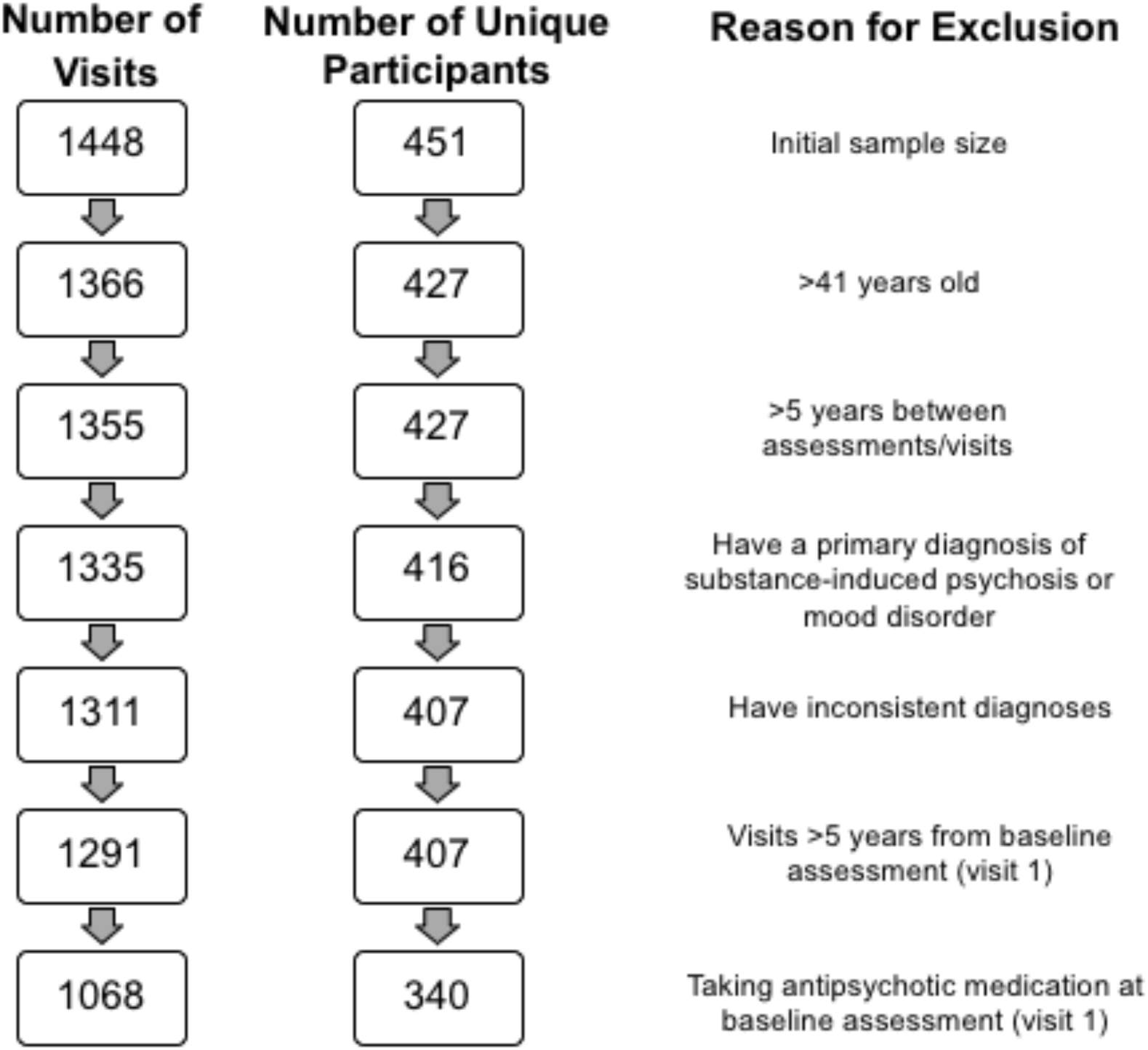
Chart of initial participant sample size, followed by exclusion criteria and number of participants remaining.

**SFigure 2:**
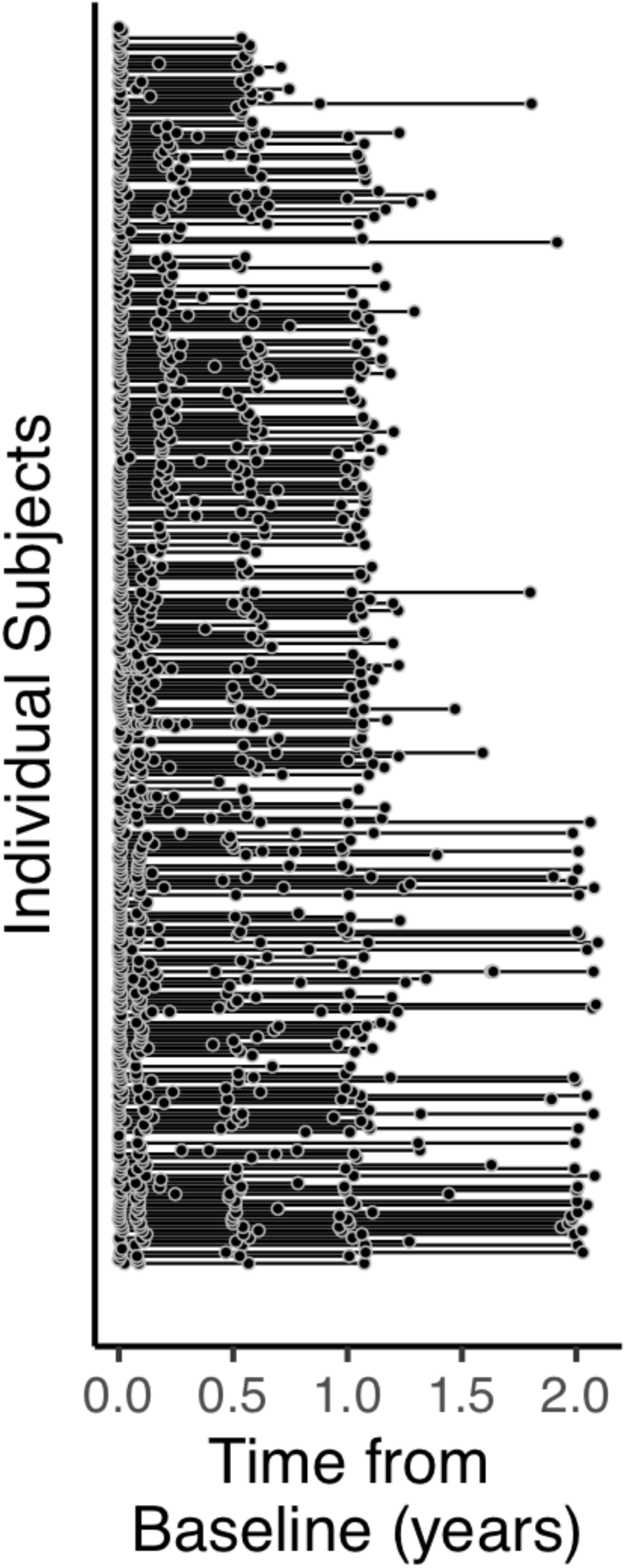
Waterfall showing length of time between visits for each participant.Individual subjects are on the y-axis and each circle represents an assessment.

**SFigure 3.**
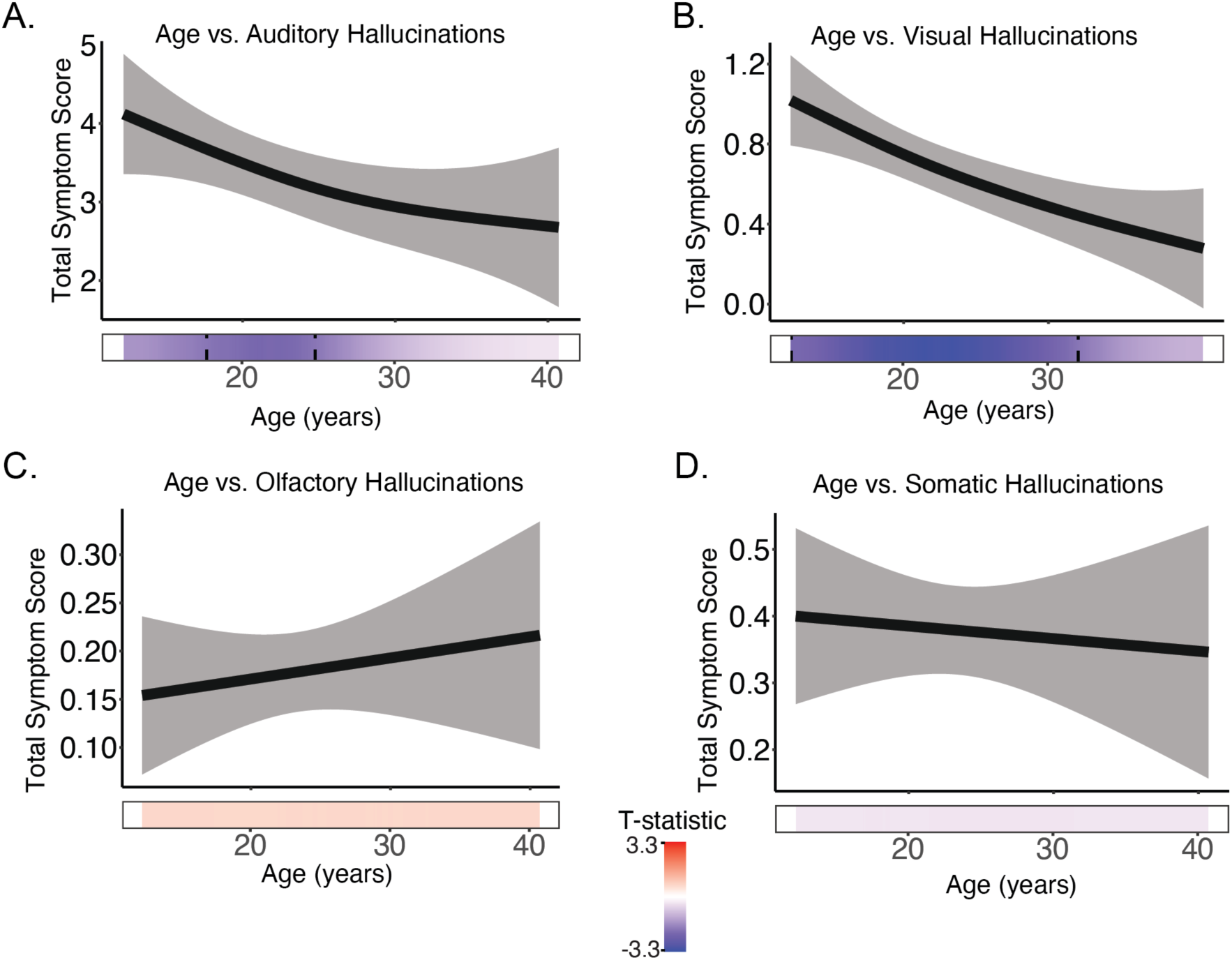
Developmental effects of A) auditory, B) visual, C) olfactory, and D) somatic hallucinations in first episode psychosis.

**SFigure 4.**
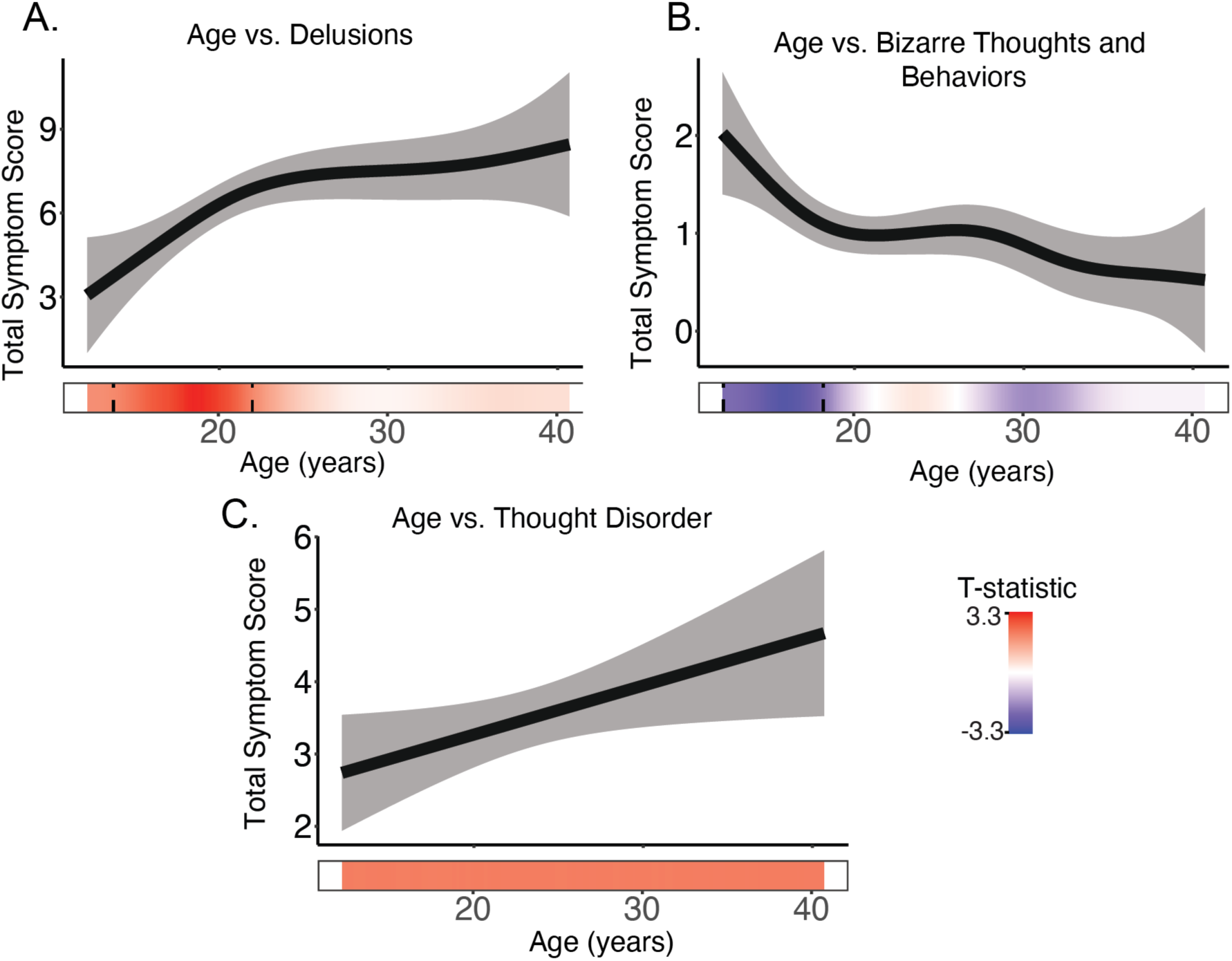
Developmental effects of A) delusions, B) bizarre thoughts and behaviors, and C) thought disorder symptoms in first episode psychosis.

**SFigure 5.**
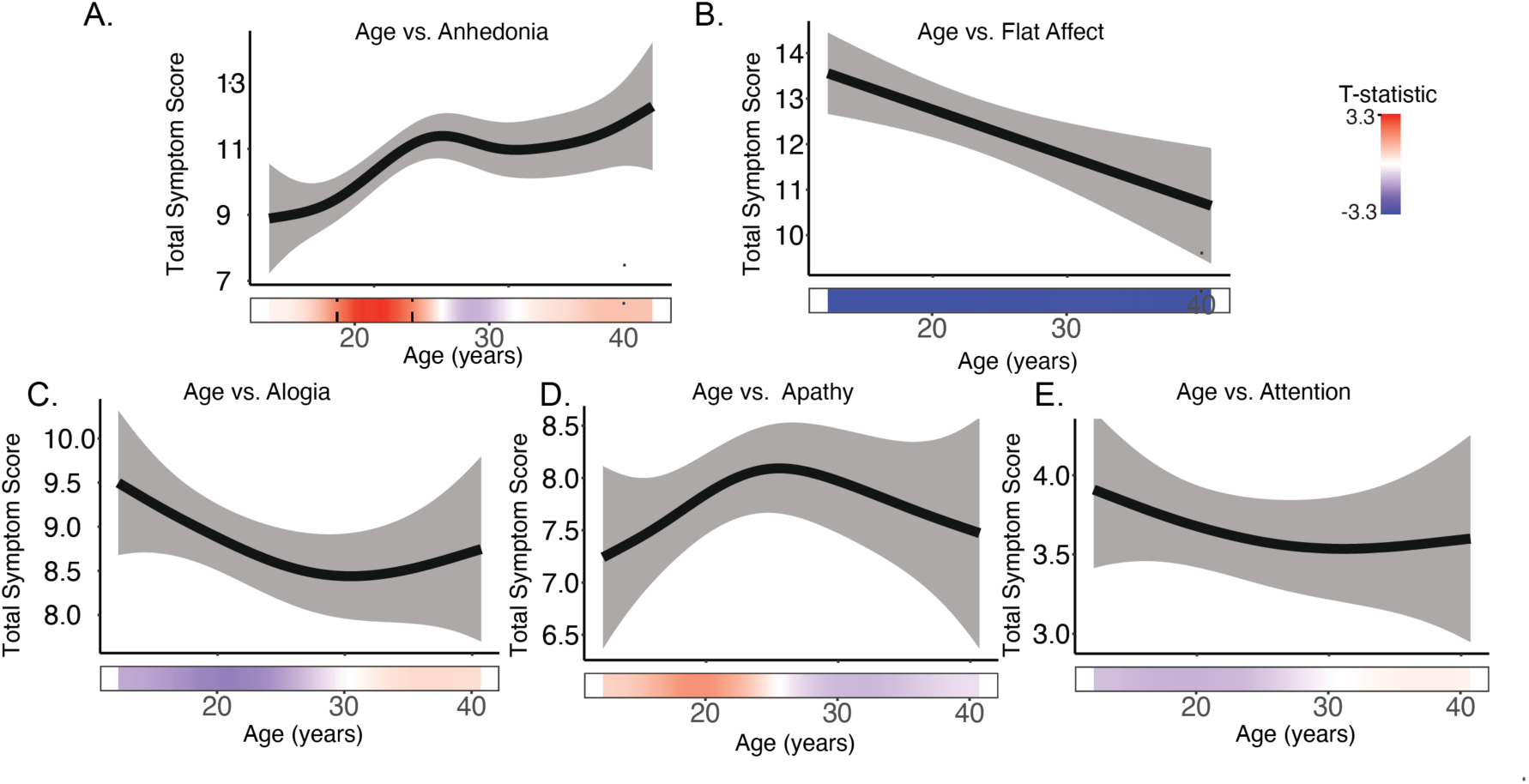
Developmental effects of A) anhedonia, B) falt affect, C) alogia, D) apathy, and E) attention in first episode psychosis.

